# Long-term stability of cellular-resolution brain-computer interface recordings after stroke

**DOI:** 10.64898/2026.08.09.742157

**Authors:** Alexander Utzschmid, Jonas Terlau, Lisa M. Held, Laura F. Schiffl, Hongbiao Chen, Göktuğ Alkan, Paolo Favero, Arthur Wagner, Jens Gempt, Bernhard Meyer, Simon N. Jacob

## Abstract

Implantable brain-computer interfaces (iBCIs) with single-neuron resolution are showing great promise for restoring mobility and communication in individuals with spinal cord injury or motor neuron disease. Stroke is the most common cause of acquired brain injury and a major contributor to long-term disability, making chronic stroke a highly relevant indication for iBCIs. However, whether stable intracortical recordings can be obtained from the structurally lesioned human brain is unknown. We report recordings from four 64-channel microelectrode arrays implanted in a participant with chronic aphasia after a large left-hemispheric stroke. The arrays targeted right-hemispheric frontoparietal regions homotopic to the damaged left-hemispheric language network. Across 111 sessions spanning 1,240 days, unit yield and signal quality remained stable. Waveform-based tracking reliably identified individual units across sessions, including across extended recording gaps. Short- and long-term unit stability was comparable to previous reports from iBCI participants without structural brain lesions, and tracked units showed consistent spiking properties across sessions. Our findings provide the first evidence that single-neuron recordings can remain stable over the long term in the stroke-lesioned human brain. They establish the feasibility of chronic, cellular-resolution iBCIs after stroke and support the development of neurorestorative applications for deficits caused by structural brain injury.

## Introduction

Implantable brain-computer interfaces (iBCIs) that record directly from the human cortex with single-neuron resolution have demonstrated remarkable potential for restoring communication, mobility and independence in neurological patients^1,2^. However, microelectrode iBCIs are currently being exclusively developed for disorders in which the brain remains structurally intact, such as spinal cord injury and motor neuron disease. Bringing microelectrode iBCI technology to patients with extensive cortical and subcortical injury remains a major unmet challenge. Stroke, traumatic brain injury and brain tumour resection, amongst others, surpass neurological conditions that preserve hemispheric brain structure in number, cost and global burden of disease by orders of magnitude^3^. Stroke, in particular, is the most common single cause of acquired brain injury and a leading cause of long-term disability^4^.

It is not known whether chronic intracortical microelectrode recordings can maintain long-term stability in the extensively lesioned brain. Importantly, global structural and functional changes following stroke may severely compromise recording reliability even at sites remote from the lesion. First, stroke-related cerebral atrophy and CSF expansion may amplify pulsatile brain movement, displacing electrodes from the surrounding neighbouring neural tissue. Second, chronic post-stroke inflammation, glial activation, and vascular remodelling extend beyond the lesion and may affect electrode-tissue interactions^5-7^. Finally, functional reorganization of remote networks to compensate for lost function, alongside functional depression of structurally intact but connected regions following stroke, termed diaschisis, can alter neural activity patterns at anatomically intact sites^8,9^. Such changes may compromise the long-term stability and interpretability of recorded signals, thereby limiting the therapeutic utility of iBCIs and challenging the justification of invasive implantation^10^.

Here, we report the first long-term investigation into the stability of cellular-resolution iBCI recordings in a patient with extensive stroke-induced cortical and subcortical structural damage. Six years prior to enrolment, the participant suffered a large left-hemispheric ischemic stroke in the middle cerebral artery territory. Clinically, the participant presented with contralateral hemiparesis and chronic non-fluent aphasia due to destruction of the perisylvian language regions. Current interventions for chronic post-stroke aphasia offer only limited functional recovery^11^, motivating an iBCI approach that bypasses the damaged language network entirely and reconstructs semantic and phonological representations from surviving neural tissue to facilitate word retrieval^12^. To this end, we implanted four 64-channel microelectrode arrays in the contralesional hemisphere, targeting cortical regions homotopic to the canonical left-hemispheric language network. To our knowledge, this is the first implantation of this kind, representing a novel iBCI-based therapeutic strategy for cognitive disorders resulting from structural lesions to the human brain^13^. Across 1240 days and 111 recording sessions, we assessed signal quality and neuronal stability in detail, a prerequisite for sustained and reliable neuroprosthetic use.

## Materials and methods

### Participant

The participant was a 51-year-old woman with a left-hemispheric ischemic stroke (M1 occlusion) six years prior to study enrolment. The stroke caused hemiparesis and non-fluent aphasia associated with extensive perisylvian infarction. Language comprehension was largely preserved. Informed consent was reaffirmed throughout the study. All procedures were conducted in accordance with the Declaration of Helsinki and approved by the TUM School of Medicine and Health ethics committee (2018-489-S-KK).

### Behavioural tasks

Six datasets comprising synchronized behavioural and neuronal data were included in this study. MonkeyLogic 2 (NIMH)^14^ was used for experimental control and behavioural data acquisition. Tasks probed language processing^15^, working memory, and mental calculation, using auditory and/or visual stimuli. The participant responded via saccades, recorded with a desktop-mounted eye tracker, or via vocal output recorded with a microphone.

### Microelectrode recordings

Four 64-channel silicon planar microelectrode arrays (“Utah arrays”, Blackrock Neurotech; 4×4 mm, 1.5 mm electrode length), totalling 256 microelectrodes and connected to a single pedestal, were chronically implanted in right middle frontal gyrus (MFG), inferior frontal gyrus (IFG), supramarginal gyrus (SMG), and angular gyrus (ANG)^16^. Extracellular signals were sampled at 30 kHz and recorded using a 256-channel Neuroport Neural Signal Processor (NSP; Blackrock Neurotech) with a Cereplex E256 headstage.

### Preprocessing and spike sorting

Raw signals were bandpass filtered (250 – 7500 Hz) and common-average referenced within each array. Spikes were sorted into units using Kilosort 1.0^17^. Cluster quality was assessed via Mahalanobis distance to the mean waveform, with a chi-squared fit used to identify outliers; spikes with a likelihood below 1/N (N = total spikes per cluster) were discarded. Units required mean firing rate ≥ 0.1 Hz and SNR ≥ 15 dB. Units were manually inspected and classified as single- or multi-unit^15^.

### Unit tracking pipeline

Units were matched based on waveform similarity, enabling cross-session tracking independent of firing rate or tuning. To this end, the unitMatch^18^ algorithm, originally developed for ultra-high-density recordings in animal models, was adapted for multielectrode arrays (MEAs). Unlike high-density probes, MEAs provide a single-channel waveform per unit, precluding the use of spatial waveform features. Instead, we leveraged the electrode geometry of the array. Given the 400 µm inter-electrode spacing, units recorded on different electrodes were assumed to originate from distinct neurons (consistent with previous studies^19,20^) and used to generate labelled training pairs.

Following spike sorting, each session was split at its midpoint, and average waveforms were computed per split for units with ≥ 500 spikes. Four similarity measures were computed across splits (“same”) and against random different-electrode units (“different”): Shape and slope similarity (Pearson correlation of waveform averages and their first derivatives), amplitude similarity

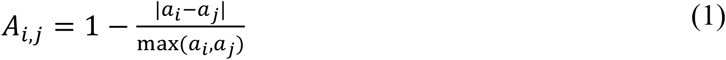

(a = trough-to-peak amplitude) and width similarity

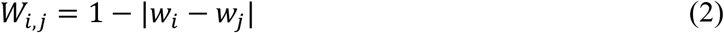

(*w*= trough-to-peak width).

Features were concatenated across sessions and min–max normalized to the interval [0 ,1]. For every electrode, an SVM trained on within-session pairs yielded decision scores for all pairwise unit combinations across sessions. Decision scores were converted to posterior probabilities via the MATLAB function fitPosterior, yielding a P(match) matrix. The upper triangle of this matrix was vectorized and sorted by descending match probability. Unit pairs were inspected iteratively and merged if the match probability of all pairwise combinations within the group of units exceeded 0.85. This followed the default UnitMatch procedure but used a more stringent matching threshold (0.85 vs. 0.5) to achieve 99 % cross-validated precision (∼1 % false matches). Merged units were assigned a unique identifier. Because all session pairs were evaluated simultaneously, units absent from intermediate sessions could still be tracked (“intermittent tracking”).

### Modelling of unit stability

For each session pair *i,j*, the proportion of units matched yielded P(track) as a function of temporal lag *k* = |*i*-*j*| days (*k* ≥ 1). Values were averaged across equal lags and fit with a biexponential decay model:

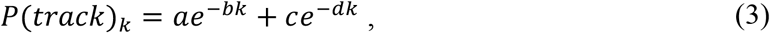

where *a, c* are the amplitudes of the short- and long-timescale decay components and *b, d* their decay constants^19^. Temporal changes in unit stability were assessed using a linear mixed-effects (LME) model with dataset ID as a random intercept and log-transformed inter-session lag and days since the first session as fixed effects, the latter visualized via partial residuals (residual + fitted contribution of this predictor).

### Functional validation

Functional stability was assessed using log-transformed inter-spike interval distributions (ISI), spike-triggered population responses (stPR) and trial-averaged responses to visual stimuli in the first dataset. Similarity was quantified using Pearson correlation. To quantify how closely tracked-pair correlations resembled true same-unit pairs, we modelled the tracked-pair distribution as a linear mixture of same- and different-neuron reference distributions,

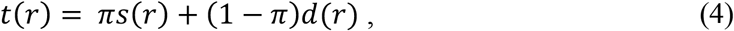

where *t(r), s(r)* and *d(r)* are the binned probability densities (50 bins) for tracked-, same-, and different-neuron correlations, and *π* is the estimated proportion of tracked pairs consistent with the same-neuron distribution. *π* was estimated via non-negative least squares, with weights normalized to sum to 1.

## Results

Across 1240 days (first recording to last recording), we acquired 111 recording sessions spanning six datasets with distinct behavioural tasks (**Fig. 1A-C**). Each dataset was acquired in blocks (up to 3 sessions per week), enabling evaluation of recording stability across both short and long timescales. Recording stability was quantified in terms of impedance, unit yield, spike amplitudes and signal-to-noise ratio (SNR). Impedances declined gradually before reaching a stable plateau (**Fig. 1D**), in line with previous reports^21^. We observed stable but regionally heterogeneous unit yields, with highest counts (single and multi-units combined) in the MFG and IFG arrays. The number of recorded units on the SMG and ANG arrays rose markedly after the first dataset (**Fig. 1E**). The overall lower yield in ANG was due to partial dislodging of the array shortly after the surgery due to cable torque, as confirmed by postoperative radiography. Median waveform amplitude was stable in IFG, SMG, and ANG arrays, with the MFG array stabilizing after an initial decline during the first dataset (**Fig. 1F**). This decrease had no effect on SNR, which remained constant across all implantation sites and recording sessions (**Fig. 1G**).

**Figure 1.**
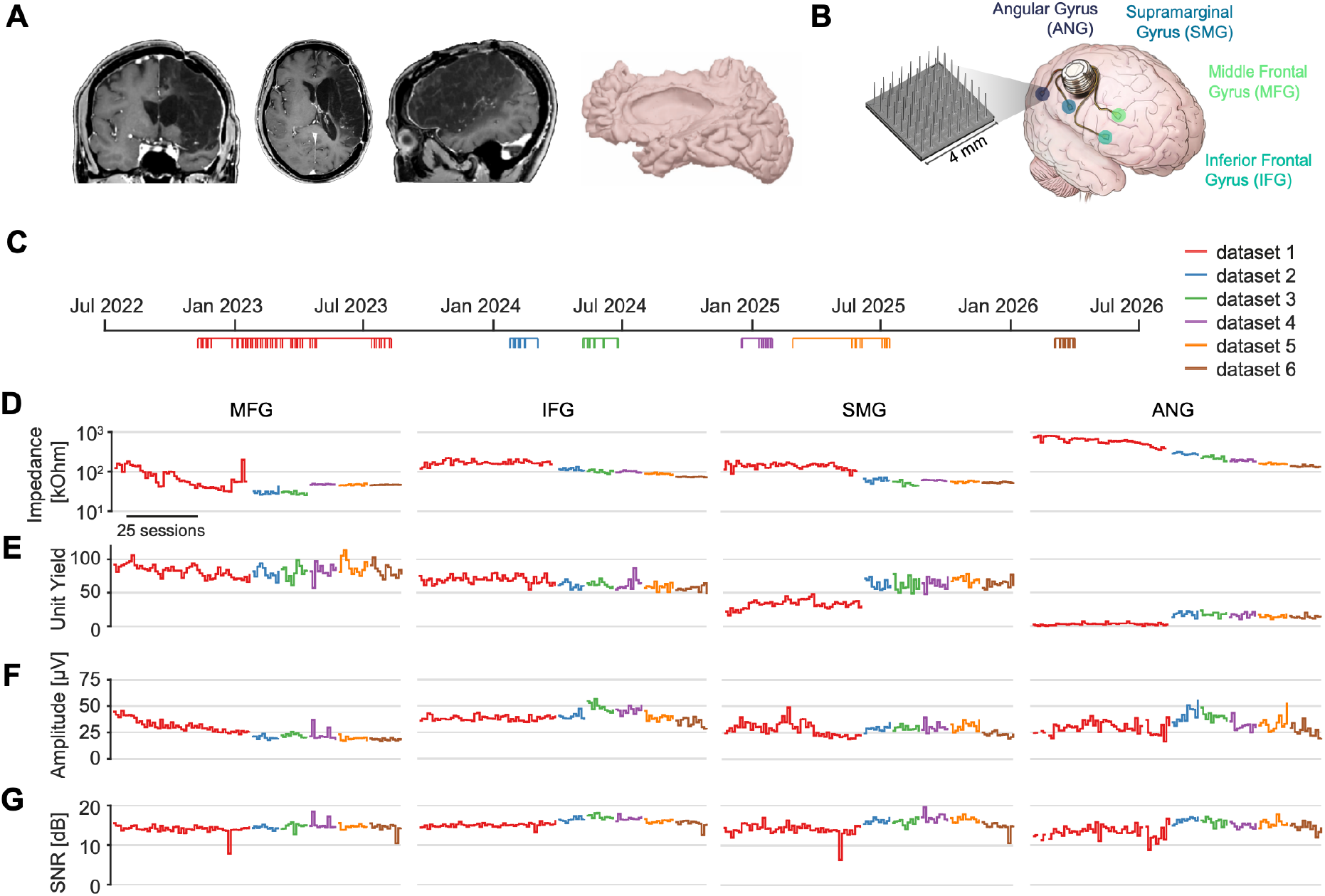
Long-term, stable microelectrode recordings in a lesioned brain. (**A**) Left: T1 MRI sections in coronal (left), axial (middle) and sagittal (right) planes showing the extent of the left-hemispheric stroke. Right: 3D reconstruction of the lesioned left hemisphere. (**B**) Schematic: Four microelectrode arrays (4×4 mm, 64 electrodes each) were implanted in the angular (ANG), supramarginal (SMG), middle frontal (MFG) and inferior frontal gyrus (IFG) of the right hemisphere. (**C**) Timeline of recording sessions. Vertical bars indicate session recording dates; colours indicate the dataset. In total, 111 sessions were recorded over 1240 days. (**D**) Unit yield (number of sorted single and multi-units per session) in MFG (83 ± 10), IFG (65 ± 8), SMG (49 ± 17) and ANG (10 ± 7, session mean ± SD). (**E**) Median waveform amplitude in MFG (26 ± 1 µV), IFG (40 ± 5 µV), SMG (28 ± 5 µV) and ANG (32 ± 8 µV, mean ± SD). (**F**) Median SNR in MFG (14 ± 1 dB), IFG (16 ± 1 dB), SMG (15 ± 2 dB) and ANG (15 ± 2 dB, mean ± SD). (**G**) Median impedance in MFG (58 ± 36 kOhm), IFG (130 ± 42 kOhm), SMG (96 ± 45 kOhm) and ANG (391 ± 225 kOhm, mean ± SD).

In addition to stable signal quality, BCI performance critically depends on the temporal stability of neuronal activity, which is constrained by longitudinal turnover in the recorded neuronal population (for example because of electrode movement). We thus tested whether individual units could be reliably tracked across multiple days. We used a waveform-based classifier (**Fig. 2A**), trained on unit pairs from session-splits (Kilosort-labelled same/different; **Fig. 2B**). Within sessions, same units were clearly identifiable based on higher similarity in waveform shape, slope, amplitude and width (**Fig. 2C**). Accordingly, the derived match probabilities P(match) were substantially higher for same versus different units (**Fig. 2D**), yielding high cross-validated classification accuracy within sessions (96.6 %). Across sessions, the classifier was applied to all pairwise combinations of units recorded on a given electrode, yielding P(match) matrices that enabled unit tracking via iterative merging of units exceeding a predefined match-probability threshold (**Fig. 2E, F**).

**Figure 2.**
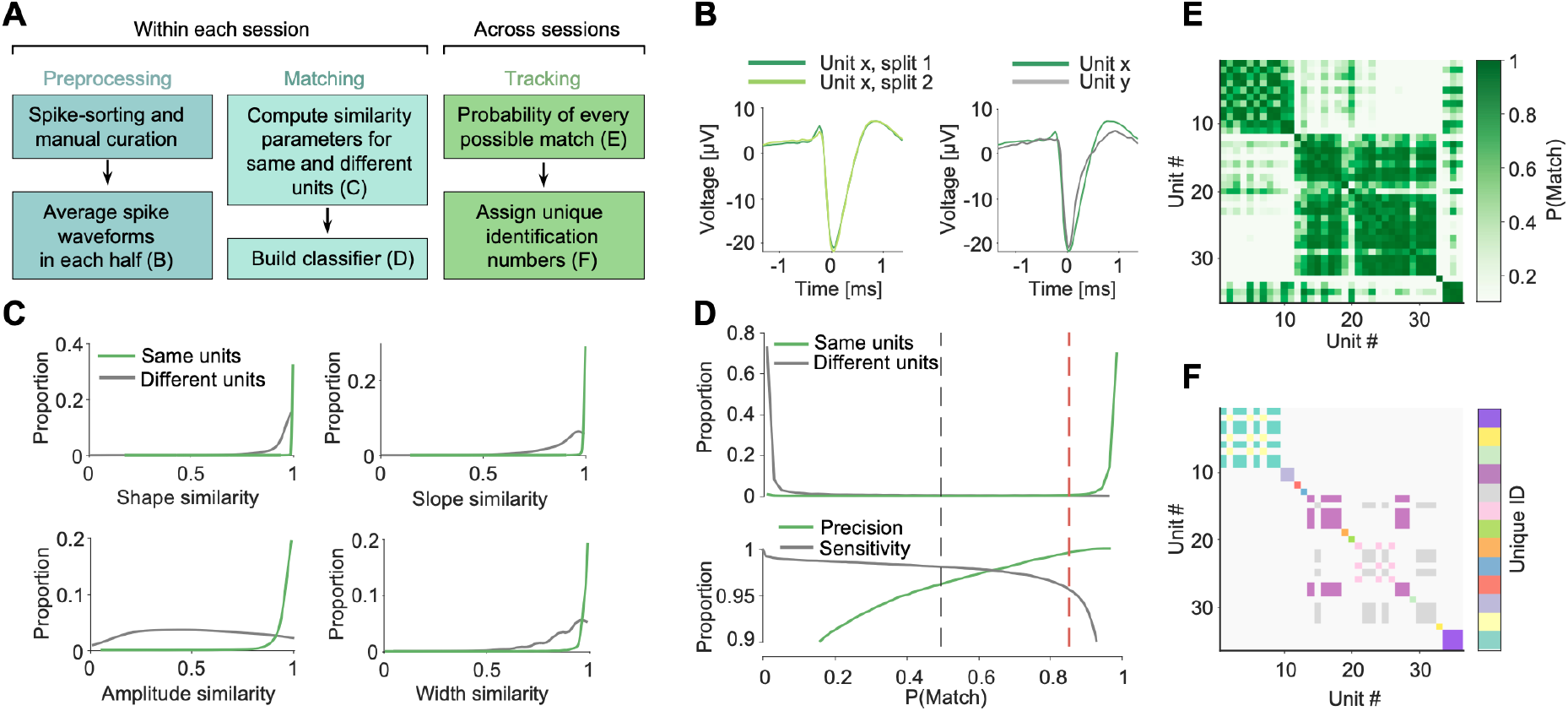
Unit tracking pipeline. (**A**) Preprocessing, matching and tracking procedures. Letters in parentheses indicate the corresponding figure panel. (**B**) Average waveform of an example unit compared across session splits (left) and to a different unit from the same session (right). (**C**) Probability density distributions of similarity features used for classification, shown for unit pairs labelled as same (green) and different (grey) within sessions. Top: Probability density distributions of match probability P(Match) for same (green) and different (grey) units. Bottom: Cross-validated classifier performance illustrating the trade-off between precision and sensitivity across decision thresholds of P(match). Dashed vertical lines indicate default (0.5, grey) and adjusted decision thresholds (0.85, red), which increased precision from 0.96 to 0.99 and decreased sensitivity from 0.98 to 0.96. Similarity matrix of all units recorded from one exemplary electrode across sessions. Units are ordered chronologically by session date. (**F**) Units are grouped into matched (i.e., identical) units when all pairwise match probabilities within a group exceed 0.85.

We compared all 111 × 111 session pairs, allowing units absent from intermediate sessions to reappear in later sessions (**Fig. 3A**, “intermittent tracking”). The mean tracking probability across pairs of consecutive sessions was 16.6 %. Individual units were tracked across temporal spans of 82.8 ± 1.8 days (mean ± SEM). We additionally applied a more conservative “continuous tracking” approach, which did not allow previously lost units to reappear in later sessions. Continuously tracked units remained identifiable across 3.2 ± 0.1 days (mean ± SEM).

**Figure 3.**
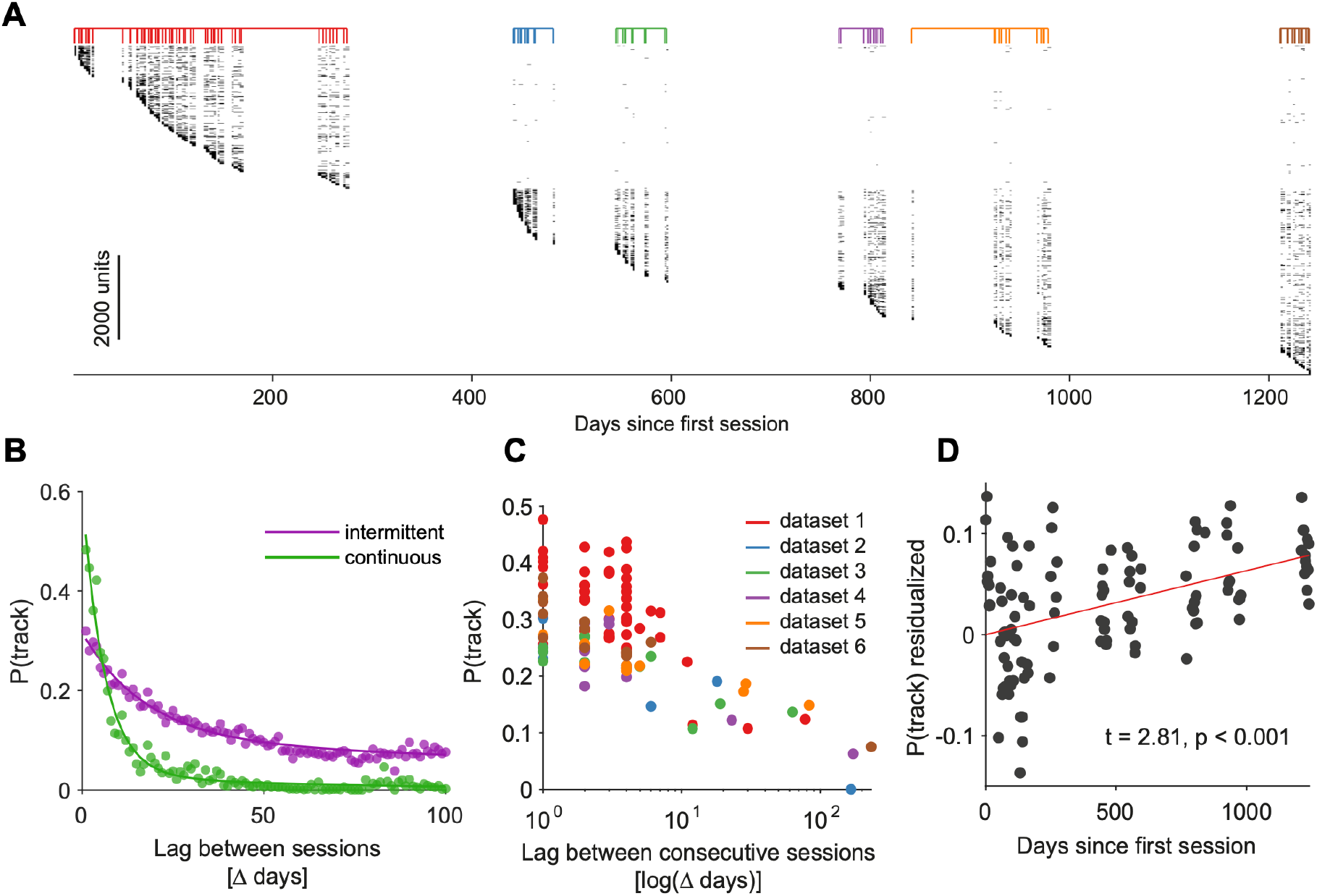
Quantification of unit stability. (**A**) Presence of unique units across recordings, intermittent tracking. Units are sorted by first appearance. Coloured vertical bars indicate the dataset. (**B**) Average P(track) as a function of days between sessions. Tracked units were divided by the number of units in the future (positive Δ days) or past (negative Δ days) session, averaged over both directions. Biexponential decay curves were fitted separately for the intermittent (purple) and continuous (green) tracking algorithms. (**C**) P(track) for consecutive sessions as a function of days between sessions, colour-coded by dataset. Each dot is one session. (**D**) Partial effect of days since first session on P(track) after accounting for session lag and dataset (t = 2.81, p < 0.001, linear mixed effects model).

To quantify short- and long-timescale components of unit stability, the proportion of tracked units across all session pairs was modelled using a biexponential decay function (**Fig. 3B**). With intermittent tracking, both the fast- and slow-decaying components closely matched those previously reported for the non-lesioned human brain^19^, exhibiting comparable amplitudes and decay constants (21.8 %, 0.06 day^−1^ and 10.2 %, 0.004 day^−1^ for the fast and slow components, respectively; previously reported: 18.7 %, 0.13 day^−1^ and 16.1 %, 0.007 day^−1^, averaged across two participants^19^).

To assess temporal changes in unit stability, we analysed P(track) between consecutive sessions across the recording period. Because P(track) decreased exponentially with increasing session lag, we log-transformed session lag to enable linear modelling (**Fig. 3C**). Specifically, we used a linear mixed-effects (LME) model to quantify the partial effect of the time elapsed since the first session (Methods). This revealed an increase in short-term unit stability, i.e., higher P(track) between consecutive sessions at later timepoints of the recording period (**Fig. 3D**).

Together, these findings show that chronic ischemic injury resulting in large structural deficits does not preclude stable long-term intracortical recordings. Unit stability was comparable to that previously reported in non-lesioned human cortex^19^ and improved over time, indicating a progressive stabilization of the electrode–tissue interface.

Finally, we tested whether tracked units showed corresponding stability in functional properties not used by the classifier, providing an independent validation of the tracking algorithm. For tracked units (**Fig. 4A**), we examined pair-wise correlations in inter-spike interval distributions (ISIs, **Fig. 4B**)^22,23^, spike-triggered population responses (stPRs, **Fig. 4C**)^24^ and responses to visual stimuli (**Fig. 4D**), compared against same- and different-unit reference pairs. Both ISIs and stPRs have been demonstrated to be highly stable neuronal characteristics^18,24^.

**Figure 4.**
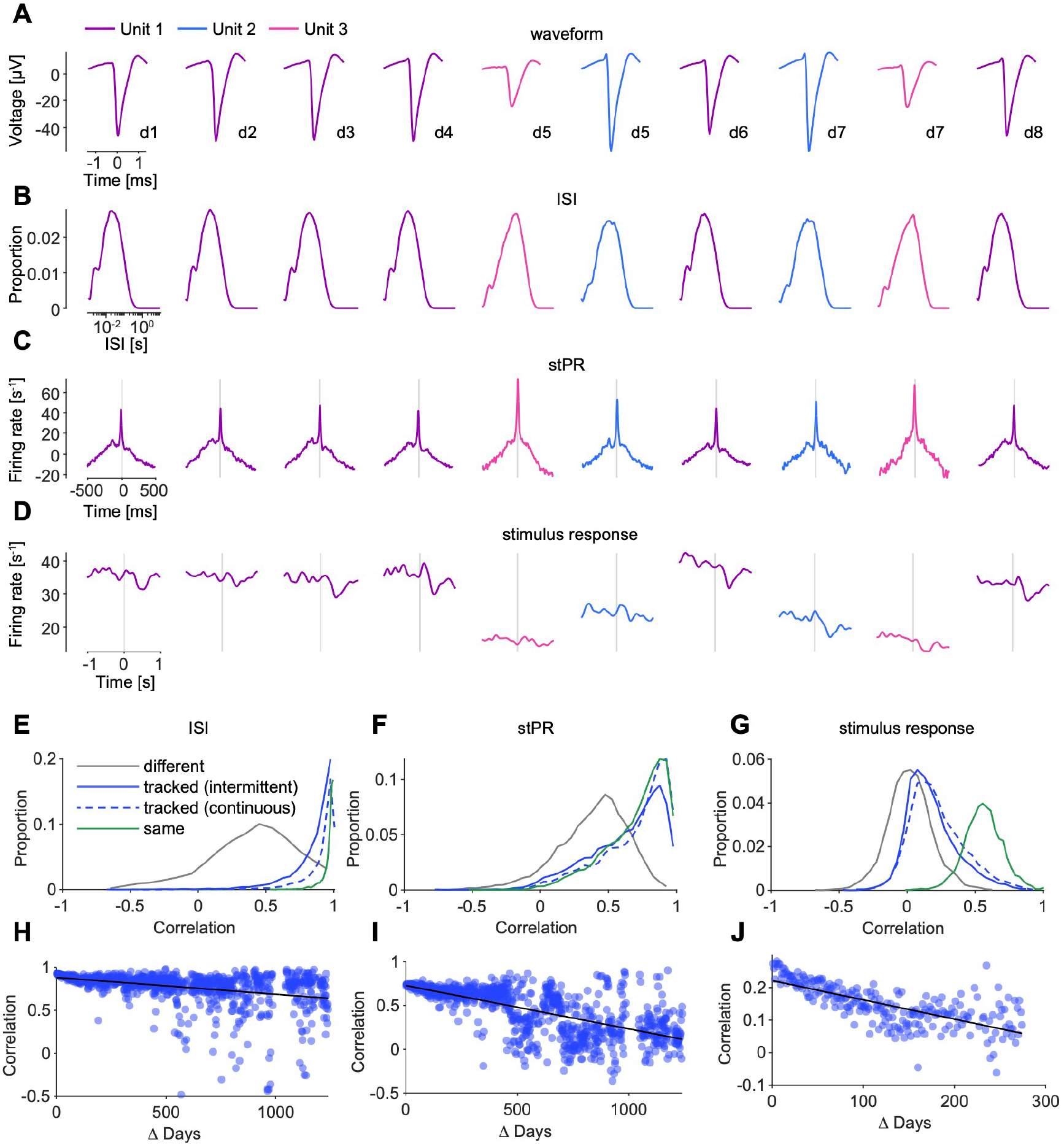
Functional stability of tracked units. (**A**-**D**) Spike waveform (A), inter-spike interval distribution (ISI; (B)), spike-triggered population response (stPR; (C)) and response to visual stimuli (D) of three example units. Intermittent tracking of Unit 1 across 8 recording sessions (d1-d8) is interrupted by the appearance of Units 2 and 3 on d5 and d7. (**E**) Distributions of the correlation in ISI distributions between same (green) or different (grey) units within days and units tracked across days using intermittent (blue, solid) or continuous (blue, dashed) tracking algorithms. 51 % (63 %) of intermittently (continuously) tracked-unit correlations were consistent with the same-unit distribution (linear mixture decomposition). (**F**) Same conventions as in (E) but for stPR. 78 % (89 %) of intermittently (continuously) tracked-unit correlations were consistent with the same-unit distribution. Same conventions as in (E) but for stimulus responses. 17 % (24 %) of intermittently (continuously) tracked-unit correlations were consistent with the same-unit distribution. (**H**) Correlation of ISI distributions between intermittently tracked units as a function of days between sessions (t = -12.29, p < 0.001, regression). (**I**) Same as but for stPR (t = -26.82, p < 0.001). (**J**) Same as (H) but for stimulus response (t = -15.81, p < 0.001). Visual stimulus responses were only analysed for the first dataset, resulting in a smaller range of inter-session intervals.

Linear mixture decomposition revealed that most tracked pairs exhibited ISI and stPR correlation profiles consistent with the same-unit distribution (**Fig. 4E, F**). Stimulus responses were less stable and more consistent with the different-unit distribution (**Fig. 4G**). To determine whether this reflected genuine changes in neuronal response dynamics, for example due to differences in the administered tasks, rather than false-positive unit matches, we examined the relationship between response similarity and inter-session interval. Because false-positive matches occur independently of inter-session interval, unit misidentification alone would not produce an interval-dependent decline in similarity. Instead, similarity in all three measures decreased significantly with increasing inter-session interval (**Fig. 4H–J**), consistent with progressive changes in the underlying neuronal dynamics. These results demonstrate consistent functional properties of tracked units, validating the waveform-based matching procedure and indicating a level of functional stability favourable for long-term iBCI performance.

## Discussion

Long-term efficacy of iBCIs depends on sustained signal quality and stability of recorded neuronal activity. Here, we demonstrate stable, high-yield single- and multi-unit recordings from the chronically lesioned human brain over nearly four years (**Fig. 1**), establishing the feasibility of chronic intracortical recordings with cellular resolution after stroke. Using a waveform-based algorithm (**Fig. 2**), we tracked individual units across extended temporal gaps (**Fig. 3**), with functional stability comparable to that observed within sessions (**Fig. 4**).

Chronically implanted microelectrodes are generally assumed vulnerable to progressive tissue encapsulation and glial scarring that can reduce signal quality^6,25^. In the post-stroke brain, persistent neuroinflammatory changes may further compromise recording stability, even at cortical sites distant from the lesion^7^, potentially amplifying the foreign body reaction to implanted iBCIs. Contrary to these concerns, the number of sorted single- and multi-units remained stable or, in SMG and ANG, even increased across the recording period. This paralleled a decline in electrode impedance to a stable plateau, consistent with the chronic electrode-tissue interface reaching steady state^21^. Speculatively, a stable glial scar may anchor electrode contacts within the neuropil, reducing micro-motion relative to local tissue. The ANG array’s transition from near-zero unit yield to consistent activity over hundreds of days exemplifies this continuing structural integration.

Beyond recording quality, long-term iBCI performance depends on the stability of neuronal activity patterns. Drift arising from multiple sources necessitates decoder recalibration and remains a major barrier to clinical adoption^26^. In the post-stroke brain, axonal sprouting, dendritic remodelling and synaptic reorganization can persist long-term and involve distributed networks beyond peri-lesional cortex^27^, potentially amplifying neuronal drift. Our results demonstrate unit stability comparable to that previously reported in participants with non-lesioned brains^19^, indicating that chronic ischemic injury does not impair longitudinal tracking. Moreover, tracked units in the contralesional hemisphere exhibited stable spiking properties across sessions, supporting the long-term stability of neuronal activity itself. These findings are particularly relevant because homotopic contralesional regions are emerging as promising iBCI targets for restoring function after stroke, including language in aphasia^15,28^ and upper-limb function in severe motor impairment^29^.

In summary, these results provide the first demonstration that stable, chronic single-neuron recordings are feasible in the lesioned human brain. Future work should test the generalizability across larger patient cohorts and determine whether similarly stable recordings can be obtained from perilesional cortex, where local tissue damage and changes in cellular and vascular physiology may more strongly influence neuronal activity and recording stability.

## Data availability

The data that support the findings from this work are available upon reasonable request from the corresponding author. Custom MATLAB code for waveform-based unit tracking is available at https://github.com/JTerlau/UnitTracking.

## Acknowledgements

Generative AI tools (ChatGPT, OpenAI; Gemini, Google; Claude, Anthropic) were used for language editing and drafting support. No AI tool was used for data analysis or interpretation. Grammarly (Grammarly Inc.) was used for language editing and grammar correction.

The authors reviewed and approved the final manuscript.

We thank our participant and her family for their pioneering spirit and extraordinary commitment, without which this work would not have been possible.

